# Probing the genomic and proteomic basis of encystment in *Oxytricha granulifera*

**DOI:** 10.1101/2025.09.17.676803

**Authors:** Miaomiao Wang, Juan Yang, Tao Hu, Zina Lin, Tian Wang, Zijia Liu, Xiao Chen, Xinpeng Fan

**Affiliations:** School of Life Sciences, East China Normal University, Shanghai 200241, China; Marine College, Shandong University, Weihai 264209, China; SDU-ANU Joint Science College, Shandong University, Weihai 264209, China

**Keywords:** cyst, mucocyst, spliceosome, transcriptomic, 6mA

## Abstract

Protozoan encystment is a crucial survival strategy for resisting environmental stresses, yet its molecular mechanisms remain poorly understood due to limited genomic resources and lack of integrated multi-omics analyses in model ciliates. In this study, we aimed to elucidate the molecular mechanisms underlying the encystment process in *Oxytricha granulifera* by sequencing and assembled its genome, and by conducting comprehensive transcriptome, proteome and morphology analyses to reveal gene expression and morphologic changes between vegetative and cyst stages. Morphological observation revealed ciliary dedifferentiation and cyst-wall formation during encystment, events respectively supported by the downregulation of microtubule dynamics related genes and the upregulation of vesicle transport related genes in cyst stage. Expanded gene families for carbohydrate metabolism, autophagy and cellular acidification align with the species’ mucocyst signature and their observed autophagic clearance, hinting at previously unrecognized mechanisms underlying cyst formation. Elevated expression of the ubiquitin-proteasome system and autophagy pathways facilitates protein turnover, and upregulation of antioxidant enzymes genes helps mitigate oxidative damage, while a rewired post-transcriptional regulation that increases spliceosome activity and alternative splicing frequency—each trend validated at the protein level. Concurrently, the methyltransferases responsible for DNA N6-adenine methylation (6mA), with homologous genes of *AMT1* and *AMT6/7* significantly downregulated. These results suggesting a multilayered regulation of ciliate encystment and the first integrated evidence that alternative splicing modulates dormancy. Our findings establish the first multi-omics framework for *O. granulifera* encystment, offering baseline data and tentative clues to the dormancy mechanism that will inform future inquiries into how single-celled eukaryotes endure adverse environments.

**IMPORTANCE:** *Oxytricha* species are widely distributed in freshwater and terrestrial ecosystems, playing significant ecological roles in microbial communities. Their ability to undergo encystment provides a powerful model for studying cellular differentiation and stress adaptation in microbial eukaryotes. This study presents the first multi-omics analysis of encystment in *Oxytricha granulifera*, revealing microbial survival strategies through enhanced protein turnover, autophagy, alternative splicing, and DNA methylation reprogramming. These findings offer fundamental insights into dormancy mechanisms and environmental adaptation in protists, advancing our understanding of microbial resilience, evolutionary innovation, and ecological success in fluctuating environments.

## INTRODUCTION

Microbial dormancy is a crucial survival strategy, allowing microbes to endure harsh conditions by entering a state of reduced metabolic activity. This phenomenon is widespread among bacteria, fungi, and protists, forming a "seed bank" that enabling microbes to disperse to new locations, and contribute to genetic variation upon resuming growth (McDonald et al., 2024; Mestre & Hofer, 2021). It also impacts microbial interactions, such as pathogens’ ability to survive in hosts and resist treatments (Gutiérrez et al., 1998; Schaap & Schilde, 2018). Microbial dormancy is characterized by reduced metabolic activity, increased stress tolerance, cessation of growth and division, and sometimes the formation of specialized structures (Corliss & Esser, 1974; Li et al., 2022). These characteristics enable microbes to survive unfavorable conditions and preserve genetic diversity. The molecular mechanisms underlying microbial dormancy involve several key processes. Regulatory proteins, such as hibernation factors, bind to essential cellular components like ribosomes and RNA polymerase, preventing their degradation and maintaining a state of metabolic stasis (Rittershaus et al., 2013). Additionally, dormancy can be triggered by the accumulation of specific signaling molecules that respond to environmental stressors (Mali et al., 2017). These mechanisms collectively allow microbes to enter and maintain a dormant state until conditions become favorable for growth and activity. Understanding the dormancy mechanisms of protists is significant for comprehending their survival strategies, evolutionary adaptability, and ecological roles (Xu et al., 2021).

Ciliates, a globally distributed group of eukaryotic microbes, have evolved an extraordinary dormancy mechanism by forming cysts to cope with diverse environmental challenges (Li et al., 2022). In recent decades, research into encystment of ciliates has focused on their drastic morphological and physiological changes, as well as underlying molecular mechanisms. Morphologically, encystment in ciliates involves several distinct stages. It typically begins with a reduction in cellular volume, followed by progressive sphericalization. The cyst wall, composed of distinct layers derived from different precursors, forms subsequently. During this process, ciliates exhibit varying degrees of ciliature resorption. Some lose both cilia and their supporting membranous structures, while others retain kinetosomes and/or microtubular structures (Grimes, 1973; Gutiérrez, 1983). During encystment, ciliates accumulate reserve grains and aggregate their mitochondria in the peripheral cytoplasmic region. The macronuclei fuse, and their chromatin condenses into small, spherical, dense bodies that are regularly scattered throughout the nuclear matrix. Additionally, various organelles within the cell undergo autophagy (Gutiérrez et al., 1998; Verni et al., 1984).

These biological events are thought to be regulated by intracellular signaling pathways that transduce environmental cues into gene expression alterations. Previous study has suggested that *Colpoda* encystment may be mediated by elevated cAMP levels resulting from adenylate cyclase activation, which is potentially regulated by the Ca^2+^/calmodulin complex (Matsuoka et al., 2009). Comparative transcriptomic analyses between the vegetative and encystment stages are increasingly being utilized to elucidate the molecular regulatory mechanisms underlying these distinct physiological phases in a growing number of ciliate species such as *Euplotes encysticus*, *Colpoda aspera*, *Pseudourostyla cristata*, and *Apodileptus* cf. *visscheri*. These studies have revealed significant pathway differences in cell cycle regulation, biosynthesis, and energy metabolism, while identifying key differentially expressed genes and signaling pathways - including cAMP, mTOR, PI3K/AKT, calcium, and AMPK—that potentially constitute a comprehensive regulatory network for encystment (Chen et al., 2018; Chen et al., 2014; Jiang et al., 2018; Pan et al., 2019; Xu et al., 2023). Pan et al. identified lncRNAs in *P. cristata* that are likely involved in encystment regulation. These lncRNAs may function by modulating their co-expressed mRNAs during encystment, potentially enhancing autophagy, protein degradation, cellular stress tolerance, and dynamic regulation of tubulin, as well as regulating intracellular calcium concentrations and inhibiting cell proliferation (Pan et al., 2021; Pan et al., 2019). Furthermore, Liu et al. (2024) proposed a 6mA-mediated epigenetic regulatory mechanism in which dynamic methylation-demethylation cycling modulates gene expression to coordinate encystment-excystment transitions in ciliates, enabling rapid environmental adaptation with minimal energy cost.

Despite significant advances in understanding ciliate encystment at the morphological and transcriptomic levels, critical gaps remain in our comprehensive understanding of this crucial survival mechanism. Currently, three major limitations hinder progress in this field: (1) The overwhelming majority of studies have focused solely on transcriptomic changes, creating a significant protein-level validation gap for putative dormancy regulators. (2) Genomic resources for encystment-capable ciliates remain remarkably scarce, with only a handful of species (e.g., *Pseudourostyla cristata*, *Colpoda illuminate*) having available genome assemblies (Jin et al., 2024; Li et al., 2024). This severe limitation prevents meaningful comparative genomic analyses to identify conserved encystment-related gene families and lineage-specific adaptations. (3) Existing studies lack integration across biological scales - while morphological changes are well-documented and some molecular pathways have been identified, the crucial links between genomic features, protein networks, and cellular restructuring remain largely unexplored.

*Oxytricha* species are widely distributed in freshwater and terrestrial ecosystems, where it thrives in a variety of ecological niches. The ability of *Oxytricha* to undergo encystment makes it an excellent model for studying cellular differentiation, gene regulation, and survival strategies in eukaryotic microbes (Lu et al., 2023). Investigating the genetic and proteomic responses underlying encystment not only reveals how microorganisms adapt to environmental stressors but also provides broader insights into eukaryotic resilience and evolutionary innovation.

In this study, we present the first high-quality genome assembly of *Oxytricha granulifera*, establishing a foundational genomic resource for this understudied ciliate. Combining transcriptomic (RNA-seq) and proteomic analyses, we systematically characterized the dynamic gene expression and protein profiles during the transition from vegetative growth to encystment. Through functional annotation of differentially expressed genes (DEGs) and proteins, we identified key molecular pathways involved in cyst formation, uncovering novel regulatory mechanisms that govern this critical survival strategy. Our multi-omics study reveals key mechanisms of encystment, advancing the understanding of stress adaptation in microbial eukaryotes.

## MATERIALS AND METHODS

### Cell culture, encystment induction and cell collection

*Oxytricha granulifera* was isolated from the soil samples collected in October 2019 from Tianmu Mountain, Zhejiang Province, China (30°19’45"N, 119°26’32"E). The vegetative cells were cultured in 500 mL sterile cell culture bottles containing Nongfu Spring mineral water, with *Chlamydomonas reinhardtii* as the sole food source, and a large number of vegetative cells (2000 cells/ml) can be obtained in 3–7 days at room temperature. For genome, transcriptome and proteome sequencing, vegetative cells were starved for 1day and purified with nylon sieves with a pore size of 1 μm (cells passed through and impurities were trapped), followed by centrifugation at 900 rpm for 5 min at 18℃ and removal of the supernatant. To induce the formation of mature cysts, the purified vegetative cells were transferred to Petri dishes and subjected to starvation conditions in culturing water for 30 days. The cysts were centrifuged at 4000 rpm for 5 minutes at 4℃, after which the supernatant was carefully removed. For transcriptome and proteome sequencing, the purified vegetative cells and cyst samples were divided into the Cyst group (cysts) and the Oxy group (vegetative cells), with three replicates per group, each containing approximately 1.5 × 10^6^ cells.

### Morphological observation by light microscope and electron microscope

Staining of encysting cells with protargol was performed to reveal the ciliature change during encystment (Wilbert, 1975). Bright field microscopy was performed with an Olympus BX 53 microscope equipped with a digital camera (Olympus DP 74).

For fluorescence microscopy, cells were fixed with Bouin’s fixative at room temperature and washed with ultrapure water in an embryo dish. Then they were stained by FITC labeled Concanavalin-A (FITC-ConA) (Alpha diagnostic International, San Antonio, USA) for 10 min and DAPI (Beyotime, China) for 1min, respectively. The cells were washed again before they were transferred onto a slide. Cells were observed in a fluorescence microscope (Nikon Ni-U, Japan) equipped with fluorescent filter blocks for ex/em 490/525 nm and 358/461 nm.

For transmission electron microscopy (Janke et al., 2005), 10-day old encysting cells were fixed for 10 minutes at 4℃ in a fixative of 2% OsO_4_ and 6.25% glutaraldehyde, then washed with 0.2M phosphate buffer and post-fixed in a fixative of 2% O_S_O_4_ and 3% potassium hexacyanoferrate for 30 minutes. The samples were washed again with phosphate buffer, dehydrated in a graded acetone series, and embedded in Epon 12 resin (Ted Pella). After ultrathin sectioning, samples were observed with a Hitachi JEM2100 transmission electron microscope at an accelerating voltage of 120 kV.

For scanning electron microscopy (Alvarez et al., 2017), samples were treated mainly according to the method described by Fan et al. (2021). Cells were fixed in Párducz’s fixative (1:4 mixture of 1% O_S_O_4_ and a saturated solution of HgCl_2_) at room temperature for 10 min. The cells were then rinsed with 0.1 M phosphate buffer, dehydrated in a graded series of ethanol, dried with a critical point dryer (Leica CPD300) and coated with gold in an ion coater (Leica ACE600). Observations were performed using a Hitachi S-4800 at an accelerating voltage of 3 kV.

### Total RNA extraction, library preparation and assembly

Total RNA was extracted using mirVana^TM^ miRNA ISOlation Kit (Ambion-1561) following the manufacturer’s instructions (Termo Fisher Scientifc, Waltham, MA, USA). The quality and concentration of the RNA samples were analyzed by NanoDrop 2000 and Agilent 2100. The mRNA was enriched using Oligo magnetic beads, and the enriched mRNA was broken into short fragments by adding interrupting agent, and the fragments were amplified into double-stranded cDNA libraries by PCR after reverse transcription with random primers. After reverse transcription with random primers, we added a splice at both ends of the fragment and amplified it into a double-stranded cDNA library by PCR.

The transcriptome data was assembled using Trinity v2.1 (parameters: --genome_guided_bam --jaccard_clip --genome_guided_max_intron 100 --full_cleanup) (Grabherr et al., 2011). The introns were identified by customized Perl scripts. The gene, CDS and transcript sequences were predicted by EuGene v1.5 (Foissac et al., 2008). For assessing the quality of the assembly, we employed BUSCO v5.4.0 (Simao et al., 2015) with the Alveolata dataset in protein mode.

### Genome and transcriptome assembly

Genomic DNA was extracted from about 1 × 10^6^ *O. granulifera* cells which were isolated from clonal cultures by using the Sodium Dodecyl Sulfate (SDS) method. The quality of the DNA was checked with a 1% agarose gel, and the purity was determined using a NanoDrop One UV-Vis spectrophotometer (Thermo Fisher Scientific, USA). The concentration of DNA was further measured using a Qubit Fluorometer (Invitrogen, USA). The extracted DNA was fragmented randomly into 350-bp fragments using Covaris ultrasonic crusher, and the ends were repaired. An adenine nucleotide was then added at the ends, and full-length adaptor sequences were connected. These libraries were purified with the AMPure XP system (Beckman Coulter, Brea, CA, USA). An Agilent 2100 Bioanalyzer and a real-time PCR were used for size distribution and quantitative analyses of the purified products. After the quality of the library was verified, all of the samples were subjected to paired-end sequencing by using the Illumina HiSeq 4000 platform with a read length of 150 base pairs (PE150).

With low-quality reads filtered using fastp (default parameters) (Chen et al., 2018), clean reads in the sequencing data were further evaluated by FastQC v0.11.9 (https://www.bioinformatics.babraham.ac.uk/projects/fastqc). The genome was assembled using SPAdes v3.14.0 (-k 21,33,55,77) (Bankevich et al., 2012). Contigs that were potential bacterial and mitochondrial contamination were removed by homologous search using BLAST v2.9.0+ (Altschul et al., 1997). Due to the low GC content typically observed in ciliates, we removed sequences from the genome with a GC content higher than 50%, as they were considered potential contaminants based on previous research (Xiong et al., 2015). In order to eliminate redundancy in the dataset, we employed CD-HIT v.4.8.1 (Fu et al., 2012) with a sequence identity threshold set at 98%. Contigs that lacked sufficient support (coverage < 5 or length < 400 bp) were subsequently excluded. We utilized QUAST v5.0.1 (Gurevich et al., 2013) with default settings to obtain statistical information about the assembled genomes. MEME (Bailey et al. 2009) was used for motif searching in the 50 and 30 subtelomeric regions. All protein-coding genes were meticulously annotated by leveraging a comprehensive suite of six renowned databases: the Non-Redundant (Mali et al., 2017) database, SwissProt, eggNOG, InterPro, Gene Ontology (GO), and the Kyoto Encyclopedia of Genes and Genomes (KEGG).

### Differentially expressed gene analysis

Hisat2 (v2.1.1) was used to map clean reads to the reference sequence using the same parameters mentioned above (Kim et al., 2019). The mapped reads for each gene loci were counted using featureCounts (v1.3.1) (Liao et al., 2014). After filtering out rDNA for gene expression analysis, Student’s t-test was employed to perform significance testing, and the results were subjected to multiple correction to obtain the adjusted P-value (q-value). The R package ’DESeq2’ (Love et al., 2014) was utilized to standardize and compare the gene expression levels between vegetative cells and dormant cysts of *O. granulifera*. Differentially expressed genes (DEGs) were defined as those with a fold change 2 and an adjusted q-value 0.1. Principal component analysis (PCA) was carried out using the princomp function of the R package ’ggplot2’ v3.4.1 (Wickham, 2016). Heat maps were plotted using the R package ’pheatmap’ v1.0.12 (https://github.com/raivokolde/pheatmap). The Gene Set Enrichment Analysis (GSEA) was adapted to be performed using a plugin in TBtools, and the plotting is carried out using R (Chen et al., 2020).

### Weighted gene co-expression network analysis

We utilized the WGCNA (Weighted Gene Co-expression Network Analysis) technique to establish a gene co-expression network, which facilitated the discovery of functional gene clusters (Langfelder & Horvath, 2008). The R package ’ggplot2’ was employed for hierarchical clustering analysis, which helped in pinpointing sample outliers. The WGCNA process involved determining the correlation of cyst samples, followed by the selection of an appropriate soft threshold power to create a scale-free network that correlated with external sample characteristics.

### Ortholog detection and phylogenetic analysis

We identified the common orthologs among the 31 species using Orthofinder v2.5.4 (-S diamond -M msa -T raxml -I 1.5) (Emms & Kelly, 2019). A maximum likelihood (ML) tree was produced with RAxML-HPC2 (Stamatakis, 2014). The ultrametric phylogenetic tree was constructed using r8s v1.81 (Sanderson, 2003), based on the rooted species tree inferred by OrthoFinder. Tree visualization was performed with MEGA v7.0.20 (Kumar et al., 2012). Significantly expanded or contracted gene families were identified using CAFE (Computational Analysis of gene Family Evolution) v5.0 with the parameter -k 5 (Mendes et al., 2021). For the estimation of divergence times, reference was made to the divergence time between *Paramecium tetraurelia* and *Tetrahymena thermophila* (median time: 609.8 MYA) retrieved from TimeTree (http://timetree.org/). 31 reference genomes were downloaded from public genome database (see supplementary Table S1). The amino acid sequences of the putative AMT family in Oxytricha granulifer were alignment with the MT-A70 protein in Tetrahymena thermophila by BLAST v2.9.0+ (Altschul et al., 1997). The conserved domains of MT-A70 family were identified by NCBI CD-search (Marchler-Bauer & Bryant, 2004). The accession numbers of genes used for phylogenetic analysis are shown in supplementary Table S2.

### 4D-label free proteomics

The samples were lysed by SDT (4% SDS, 100mM Tris-HCl, 1mM DTT, pH7.6) buffer for protein extraction. The amount of protein was quantified with the BCA Protein Assay Kit (Bio-Rad, USA). The protein suspensions were digested with trypsin overnight at 37 ℃. The peptides of each sample were desalted on C18 Cartridges (Empore™ SPE Cartridges C18 (standard density), bed I.D. 7 mm, volume 3 ml, Sigma), concentrated by vacuum centrifugation and reconstituted in 40 µL of 0.1% (v/v) formic acid. Samples were separated by using Easy-nLC system. Buffer A is 0.1% formic acid and Buffer B is 84% acetonitrile and 0.1% Formic acid. Samples were loaded onto a reverse phase trap column (Thermo Scientific Acclaim PepMap100, 100 μm × 2 cm, nanoViper C18) and separated with the C18-reversed phase analytical column (Thermo Scientific Easy Column, 10 cm long, 75 μm inner diameter, 3 μm resin) at a flow rate of 300 nl/min controlled by IntelliFlow technology. LC-MS/MS analysis was performed on a timsTOF Pro mass spectrometer (Bruker, Germany). The mass spectrometer was operated in positive ion mode. The ion source voltage was set at 1.50 kV and the peptide parent ions and their secondary fragments were detected and analyzed using a high-resolution TOF. The mass spectrometry scan range was set to 100–1700 m/z. Data acquisition was conducted using the parallel accumulation serial fragmentation (PASEF) mode. The 10-fold PASEF mode acquisition of mother ions was performed after first-level mass spectrum acquisition, using a cycle window time of 1.17 s. For secondary spectra with charge numbers of 0 to 5, the dynamic exclusion was set at 24 s. The MS raw data for each sample were combined and searched using the MaxQuant 1.6.14 software for identification and quantitation analysis.

### Gene expression validation by RT-qPCR

The reaction of qRT-PCR was performed with a QuantStudio 5 instrument (Thermo Fisher Scientific, USA). For each sample, reactions were performed in three independent wells. The PCR system was 20 µL in total and contained 10 µL of 2 × ChamQ Universal SYBR qPCR Master Mix (Vazyme, Q711-02), 0.4 µL of upstream and downstream primers (10 µmol/L), 1 μL of cDNA template, and ddH_2_O to the final volume. The reaction procedures were as follows: (i) holding stage, predenaturation at 95 ℃ for 30 s; (ii) PCR stage, 40 cycles were performed at 95 ℃ for 3 s and 60 ℃ for 30 s; (iii) melt curve stage, 95 ℃ for 15 s, 60 ℃ for 60 s, and 95 ℃ for 15 s. The primer sequences and gene information are listed in supplementary Table S3. All of the reactions were repeated in triplicate and the relative expression levels were calculated using the 2^−ΔΔCt^ method (Adams, 2020).

## RESULTS

### Morphological transition of *Oxytricha granulifera* from the vegetative cells to cysts

Cells in the vegetative stage of *Oxytricha granulifera* were long ellipsoidal in shape and measured 80-120 μm long and 30-50 μm wide, with two large macronuclei and two or three micronuclei (Fig. 1A, C, E). Mucocysts were spherical in shape and about 0.5 μm in diameter, and distributed on both dorsal and ventral surface in a longitudinal and irregular arrangement; there were 4,500-5,000 mucocysts in each individual (n = 3); their positive staining with FITC-conA indicates their glycoprotein component since Con A, binds to internal or non-reducing terminal α-mannosyl residues (Sandoval-Altamirano et al., 2017) (Fig. 1D, E). The adoral zone of membranelles (AZM) occupied about one-third of body length. Frontal-ventral-transverse cirri arranged in the typical 8-5-5 pattern. Left and right marginal rows were gently curved at posterior end, and composed of 23-27 and right 23-31 cirri respectively (Fig. 1C; Fig. S1A). In the TEM sections, most mucocysts were beneath the pellicle, and they were elliptical in shape with an electron-light cavity in anterior part; mitochondria and starch granules were irregularly dispersed in the cytoplasm (Fig. 1H, I).

**Figure 1.**
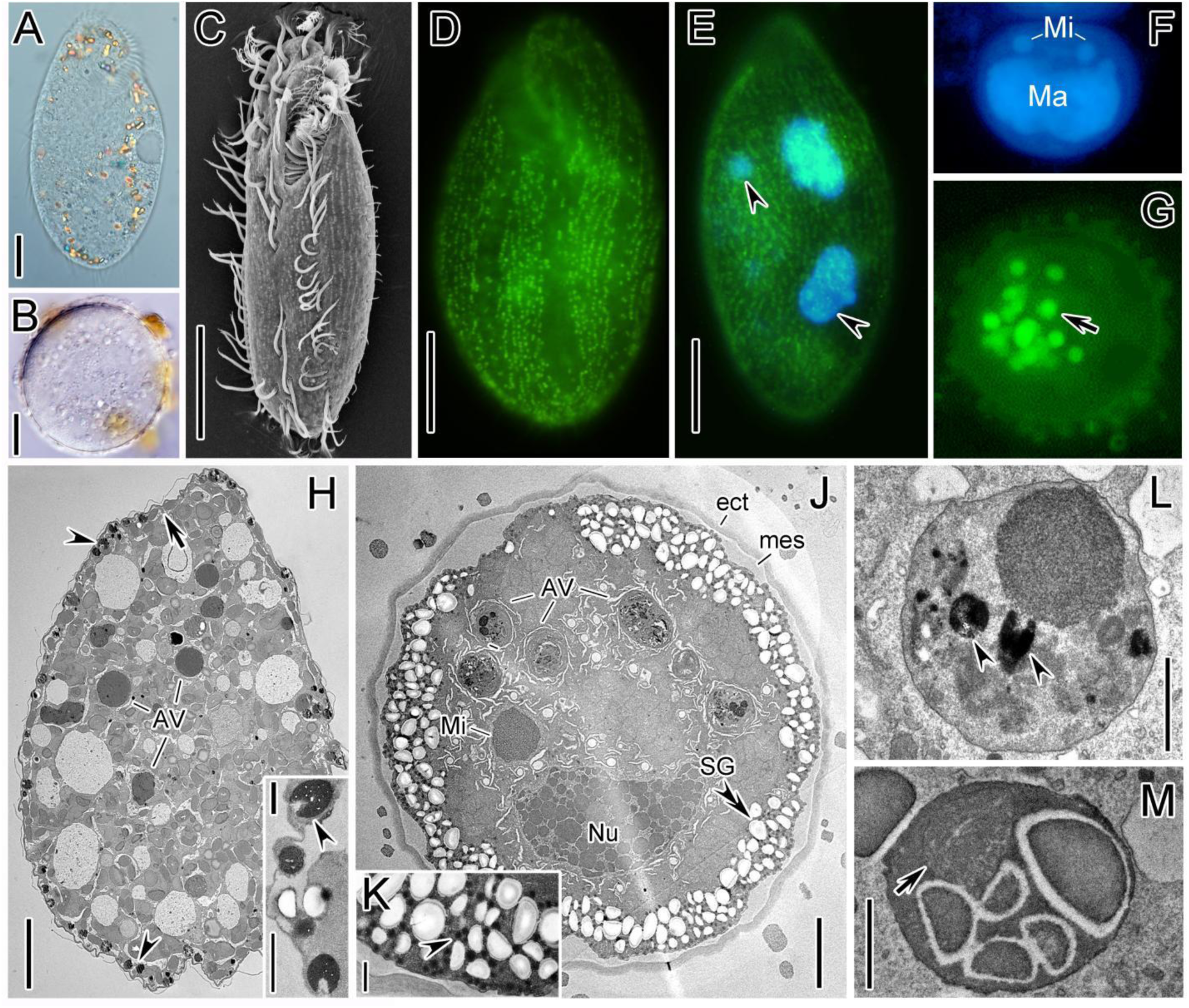
Major morphological changes during encystment of *Oxytricha granulifera*. (A, B) Vegetative and cyst stages in vivo. (C) SEM image of vegetative cell. (D, E) FITC-ConA fluorescence staining and DAPI fluorescence staining of the vegetative cell, showing the longitudinally arranged mucocysts distributed on both sides (green) and two macronuclei (blue), arrowhead mark the micronuclei which usually accompanied with each macronucleus. (F, G) Mature cyst stained with DAPI (F) and FITC-ConA (G), showing the fused macronucleus (Ma), and the autophagic vacuoles (arrow) that enclose mucocysts. (H, I) Encysting cells without cyst wall, showing the mitochondria (arrow) and mucocysts (arrowheads) beneath the pellicle. (J‒M) Encysting cells with cyst wall, showing a large number of granulars (likely the cyst wall precursors) (arrowhead) and starch granules (double-arrowhead) near in the cortex, the ectocyst (ect) and mesocyst (mes) of the cyst wall, the nucleoli of the fused macronuclei, and the homologous nucleoli (Mi), and the autophagic vacuoles enclose mucocysts (arrowheads) and mitochondria (arrow), the endoplasmic reticulum around the autophagic vacuoles (double-arrowhead). AV, autophagy vacuoles; ect, ectocyst; Ma, macronucleus; mes, mesocyst; Mi, micronucleus; Nu, Nucleolus; SG, starch granule. Scale bars: 10 μm (A‒B), 20 μm (C‒E), 5 μm (H, J), 1 μm (I, K, L, M).

In the early stage of encystment, the body shortened posteriorly, both of the marginal rows became distinctively curved, and the transverse cirri and possibly the pre-transverse ventral cirri dedifferentiation; and the proportion of the AZM to body length increased but the cirri in the frontal area did not change (Fig. S1B). Later, the body shape was more oval (AZM occupied about 2/3 of body length), the cirri in left and right marginal rows were decrease to about half of their original number because the dedifferentiation from posterior to front; transverse cirri and pre-transverse ventral cirri disappeared completely, and postoral ventral cirri began to dedifferentiate (Fig. S1C). The de-differentiation of buccal apparatus began with the fragment of AZM, i.e., the membranelles in posterior part separated from adoral zone and de-differentiated (Fig. S1D, E). At this time, the posterior frontal-ventral cirri disappeared; and meanwhile, the left marginal row completely disappeared and the right marginal row contained only several cirri (Fig. S1E).

Mature cysts were spherical and 30-50 μm in diameter, containing FITC-conA positive vesicles (likely mucocysts in autophagic vacuole) (Fig. 1B, G). In TEM sections, there were two cyst wall layers in the immature cyst: the outer wall layer was highly electron-dense with the thickness of 0.4-0.6 μm, while the inner layer was electron-lucent and uneven with the thickness of 0.2-2 μm (Fig. 1J). The nucleoli in the fused macronucleus were more homogeneous and arranged more tightly compared to those of encysting cell (Fig. 1F, J). No kinetosomes or cilia remained in the cell cortex; while the starch granules and spherical electron-dense granules densely arranged near in the cortex (0.1‒0.2 μm in diameter) (Fig. 1J, K). A few autophagy vesicles containing mucocysts, starch granules and mitochondria were also observed (Fig. 1L, M).

### Genomic features and evolutionary analysis in *Oxytricha granulifera*

Paired-end reads sequencing data (15 Gb in total) were acquired from *O. granulifera* macronuclear genomic DNA and the genome was assembled using SPAdes (Fig. 2A). After filtering out the bacteria and mitochondria related sequences, the genome assembly was de-redundified with the sequence similarity cutoff of 98%, and poorly supported contigs (coverage < 5 or length < 400 bp) were discarded. The final information of the genome was evaluated by QUAST (Fig. 2B). This assembly comprised 16,944 contigs (41.7 Mb) with the N50 of 3,112 bp. The chromosomes of *O. granulifera* are approximately 2000 bp in length, with most contigs containing one or two genes, which are characteristic features of typical nanochromosomes a characteristic genomic architecture shared with other spirotrichs (*Oxytricha trifallax*, *Euplotes vannus*, *Strombidium stylifer*) (Fig. 2B-D). Nanochromosomes usually carry single genes and have conserved potential cis-regulatory elements (CREs) in the 5’ subtelomeric regions, which regulate gene expression independently of each other (Zheng et al., 2021). This nanochromosomal organization in the macronucleus provides critical adaptive advantages, as it enables rapid gene expression regulation in response to environmental changes, thereby enhancing cellular function and survival in ciliates (Chen et al., 2014; Swart et al., 2013). The GC content of the genome was 33.43% (Fig. 2B). Gene sizes were concentrated between 2,000-3,000 bp and 22,248 genes were predicted by eggNOG (Fig. 2B). Similar to other spirotrichs, *O. granulifera* also has a higher GC content (the GC content of other ciliates is approximately 30%), a higher gene density, lower repetitive sequences, and a higher recombination rate (Chen et al., 2019; Li et al., 2021; Swart et al., 2013). The telomeres of *O. granulifera* had the sequence motif of [C_4_A_4_]_n_ (Fig. 2E). A canonical boundary motif of ’GT-AT’ was present in most predicted gene introns, which were short in size (20-30 nt) (Fig. 2F), and most genes had one exon (Fig. 2G). Genome integrity was assessed by BUSCO, with 86.5% complete and 5.3% fragmented (Fig. 2H). Ciliate, often employ the standard stop codons in unconventional manners, diverging from the typical usage observed in other eukaryotic organisms (Lozupone et al., 2001). In *O. granulifera*, UGA was mostly used as a stop codon, as is the case with other spirotrich ciliates (Fig. 2I).

**Figure 2.**
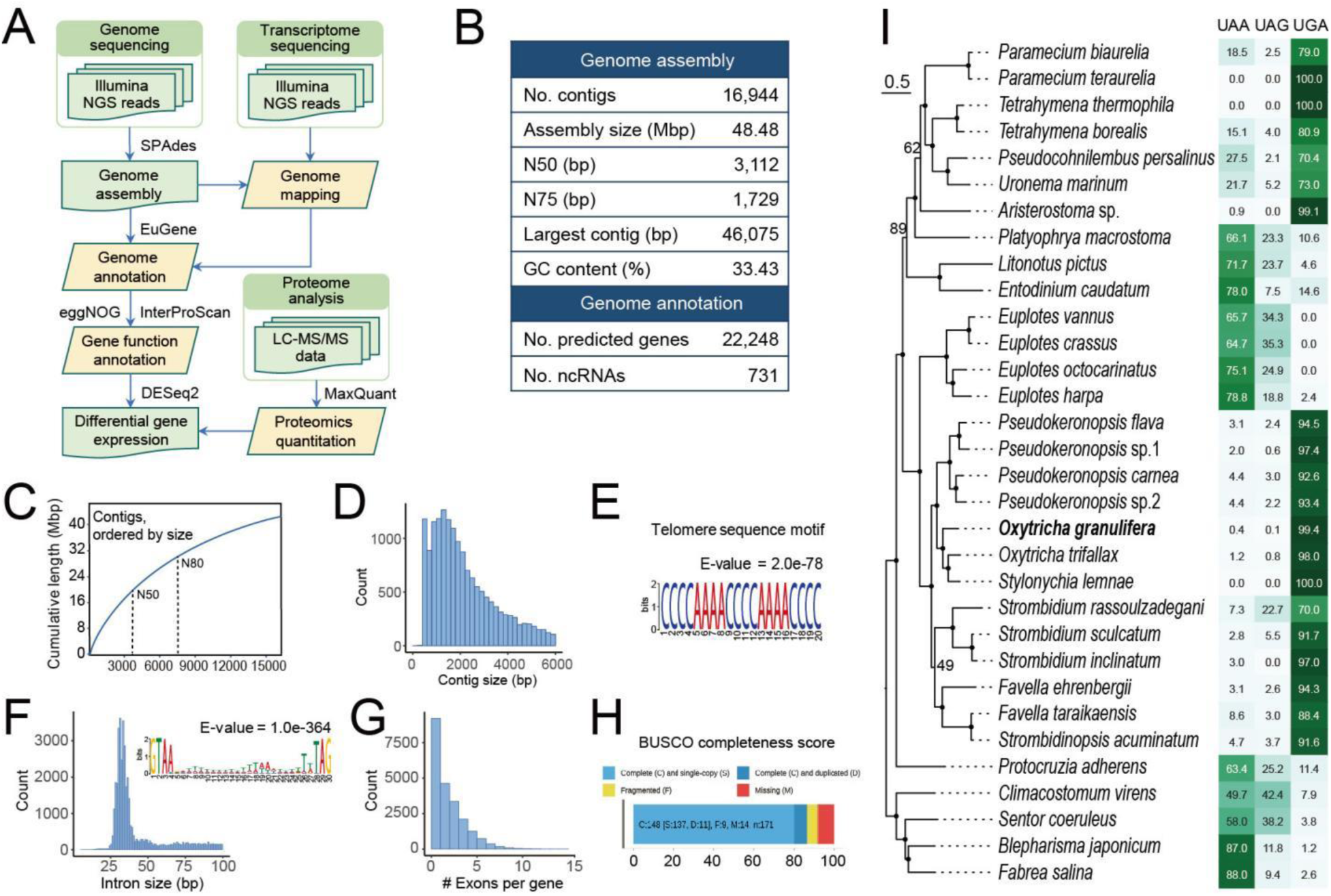
Sequencing and assembly features of somatic genome of *Oxytricha granulifera*. (A) Schematic diagram of the strategy for the genome assembly and annotation. (B) Genome assembly features. (C) Distribution of cumulative length of the genome assembly. The N50 value represents the contig length at which 50% of the assembled genome is covered by contigs of this length or longer, while the N80 value indicates the contig length covering 80% of the genome. (D) Contig size distribution across the genome assembly. (E) Potential telomere sequence motif of the genome assembly. (F) Distribution of intron sizes and the sequence motif of the intron between 20 and 30 bp. (G) Exon distribution within genes. (H) BUSCO assessment of genome assembly completeness. The results are categorized as complete (C), fragmented (F), and missing (M) BUSCOs, providing a quantitative measure of how many of the expected core genes are fully present, partially represented, or absent from the assembly, respectively. (I) Phylogenetic tree based on orthofinder by maximum likelihood (ML) method. The scale bar corresponds to 30 substitutions per 100 nucleotide positions. The adjacent heatmap to the right of the phylogenetic tree illustrates the utilization of stop codons across all 32 species ciliates.

The speciation times of *Oxytricha granulifera* and 31 other ciliates (involving Oligohymenophorea, Colpodea, Litostomatea, Spirotrichea and Heterotrichea) were inferred based on the phylogenetic analysis of amino acid sequences (Fig. 3A). Phylogenomic analyses have shown that *O. granulifera*, the type species of *Oxytricha*, is basal to the clade containing *O. trifallax* and *Stylonychia lemnae*. The divergence of *O. granulifera* occurred earlier (∼ 400 mya) than the other two members of Sporadotrichida, *O. trifallax* and *S. lemnae* which diverged from each other at ∼ 250 mya. *O. trifallax* accompanied by a large-scale gene family expansion (2895 gene families). The GO enrichment analysis showed that a total of 155 GO terms were significantly enriched (p < 0.05) in *O. granulifera*, among which the top 20 terms after ranking by q-values were collected including each aspect (molecular functions, biological processes, and cellular components) (Fig. 3B). They were predominantly associated with membrane, transmembrane transport, peptidase activity, proteolysis and carbohydrate metabolism, which confers *O. granulifera* with enhanced adaptive capabilities in nutrient acquisition, energy homeostasis and environmental stress resistance. The extensive presence of mucopolysaccharides or glycoproteins characterizing as α-mannose in the mucocysts distributed across the cortex of *O. granulifera* potentially correlates with the gene expansion (particularly mannose metabolic process, hydrolase activity) detected in this species. Specifically, the mannose metabolic pathway provides sugar precursors for synthesizing extrusomal mucopolysaccharides/glycoproteins, while O-glycosyl hydrolases remodel their structures through selective cleavage of α-mannoside linkages (Lehle et al., 2006; Spiro, 2002). In contrast, neither *O. trifallax* nor *S. lemnae* possess mucocystsand they are phylogenetically assigned to the subfamily Stylonychinae. These findings provide a genomic reference for the true *Oxytricha* and collectively support the systematic hypotheses that *O. trifallax* is not a member of *Oxytricha*, but rather belongs to the subfamily Stylonychinae (Foissner & Berger, 1999; Lu et al., 2025; Wang et al., 2025).

**Figure 3.**
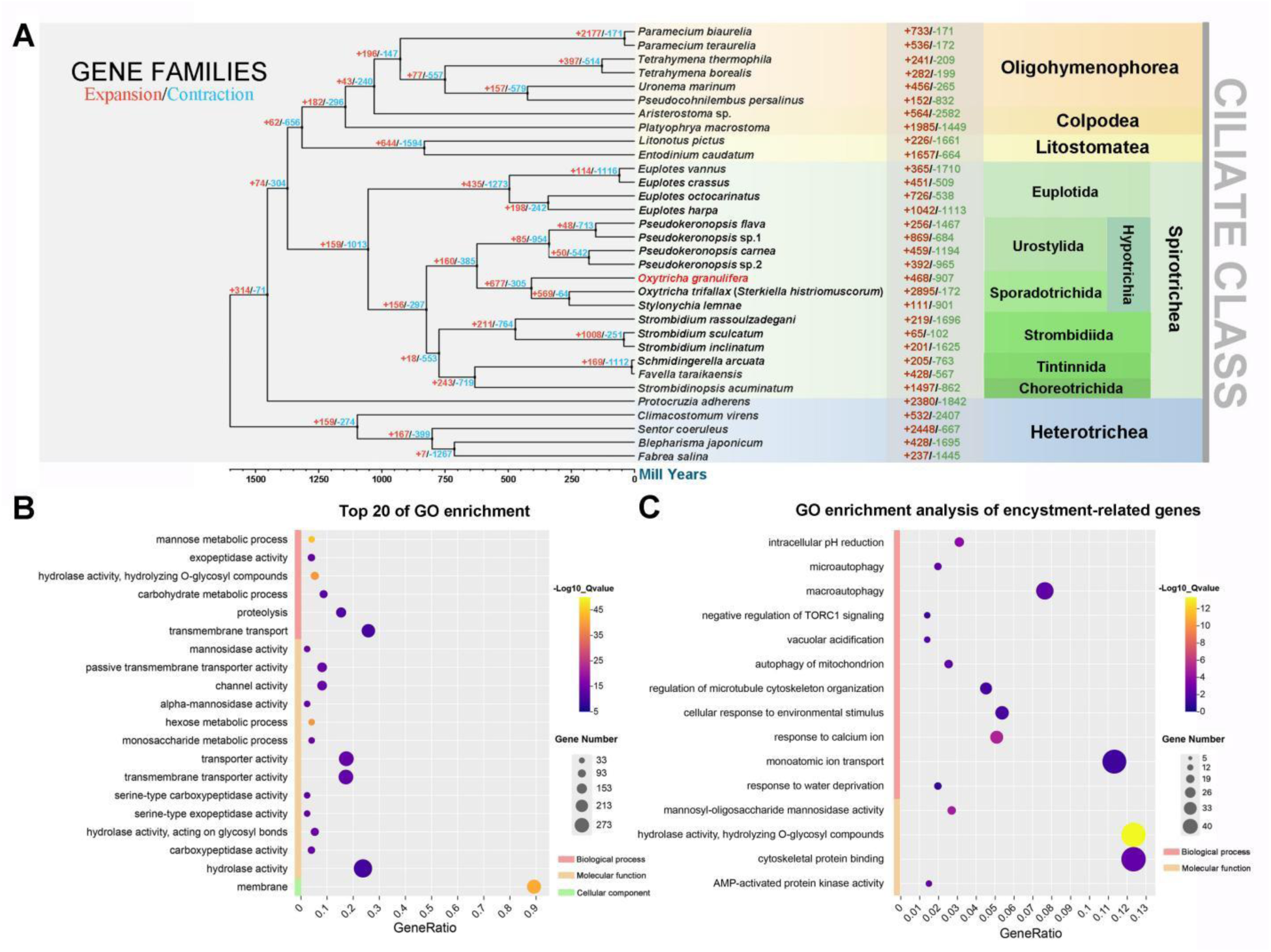
Analyses of evolutionary history and gene family expansion and contraction of *Oxytricha granulifera*. (A) The phylogenomic tree, divergence times, and gene family expansion and contraction for *O. granulifera* and 31 other species. The numbers at nodes indicate the number of gene families expanded and contracted at different evolutionary time points. Numbers following species names represent expanded and contracted gene families for that species. (B, C) The top 20 GO terms (B) and encystment-related GO terms (C) in significantly expanded gene families. GO, Gene Ontology; GeneRatio, the ratio of enriched gene number to all gene number in this pathway term.

Additionally, we identified potential multiple encystment-associated terms among the significantly enriched GO terms in expanded gene families of *O. granulifera* (Fig. 3C) according to previous studies of ciliate gene family expansion, encystment mechanisms, and encystment-related transcriptomes (e.g., Jin et al., 2023, 2024; Pan et al., 2019, 2021).

### Transcriptome analysis reveals the genetic basis of morphological changes during vegetative-cyst transition

After normalizing the read counts, the similarity between samples from different tissue types was evaluated by generating a principal component analysis (PCA) plot using the normalized count data. PCA of the dataset showed that the samples segregated primarily based on the cyst stage (Fig. 4A). Transcriptome profiles showed a dramatic change in cyst stage cells, compared to those in vegetative stage (Fig. 4B). To identify differentially expressed genes (DEGs), we plotted the volcano plot for each transcript (Fold change > 2, q-value < 0.1) and found 9,368 DEGs (7,032 downregulated genes and 2,336 upregulated genes, Fig. 4C). To validate the results of transcriptome analyses, five genes were selected to confirm the expression level using RT-qPCR. The fold change values derived from RT-qPCR and RNA-seq expression data were predominantly in agreement (supplementary Fig. S2D).

**Figure 4.**
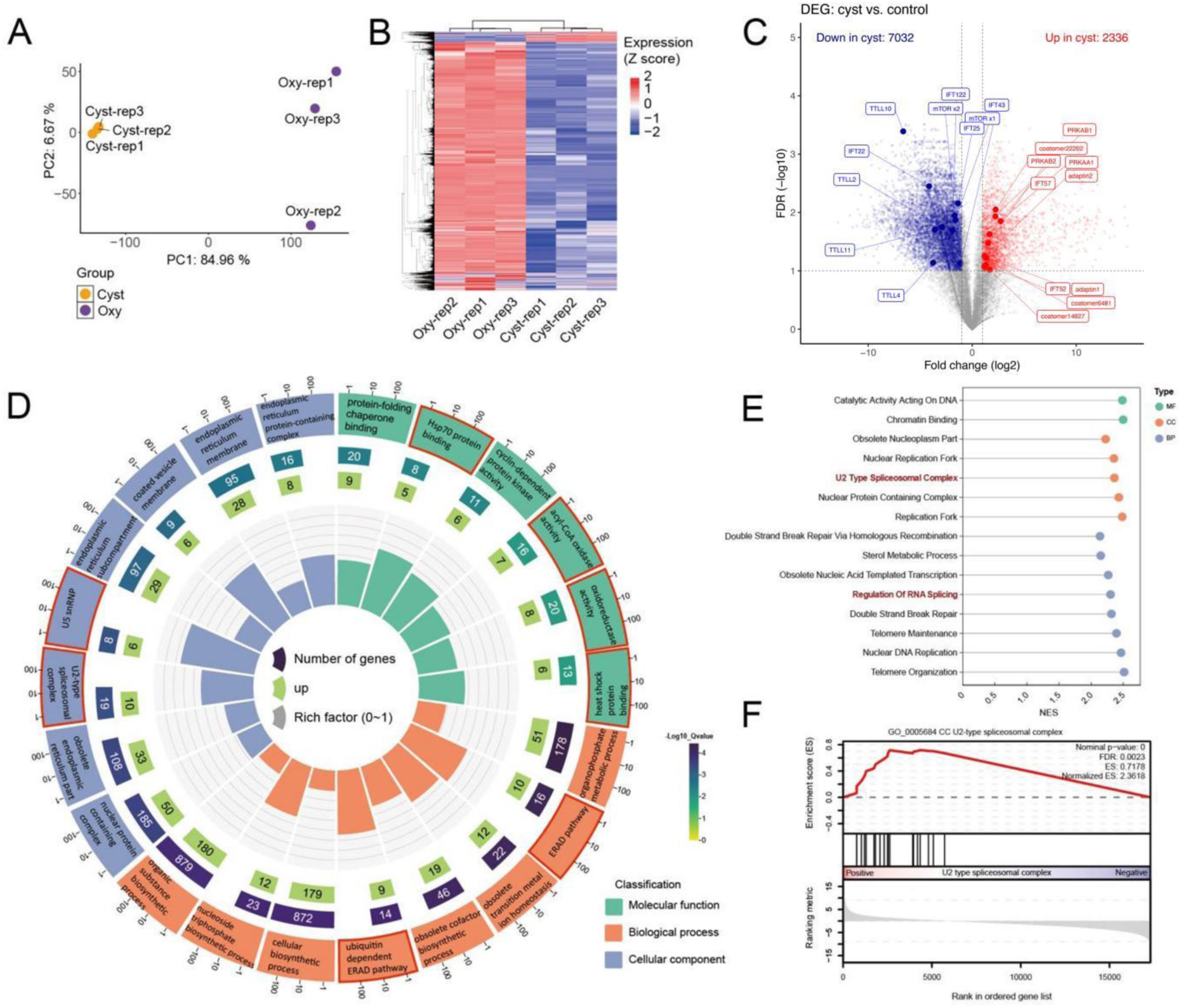
Gene expression changes between vegetative and cyst stages of *Oxytricha granulifera*. (A) Principal component analysis (PCA) based on gene expression profiles. Each point represents a sample. Orange dots represent the vegetative stage (Oxy), while purple dots represent the cyst stage (Cyst). (B) Heatmap of gene expression across different stages. (C) Volcano plot of gene expression. with fold-change thresholds of 2 and an adjusted P-value threshold of 0.1. (D) GO Circle plot. GO circles plot shows the enriched of molecular functions, biological processes and cellular components. The inner ring is a bar plot where the height of the bar indicates the enrich factor of the term, the color indicates GO terms that the corresponding genes are associated with. The outer ring displays the upregulated genes in each term. The third ring shows the total number of genes associated in this term and color corresponds to the significance (-log10 P-value). The outermost ring displays the names of the GO term. The enriched term associated with encystation are highlighted in red. (E) Result of Gene Set Enrichment Analysis (GSEA). Normalized enrichment scores were shown in x-axis. The y-axis lists the enriched biological processes and molecular functions, categorized by Gene Ontology (GO) annotations into molecular function (MF, green), cellular component (CC, orange), and biological process (BP, blue). (F) GSEA analysis of U2-type spliceosomal complex.

We performed GO enrichment analysis for the DEGs, the upregulated genes were enriched in U5 snRNP, U2-type spliceosomal complex, ubiquitin-dependent ERAD pathway, heat shock protein binding, oxidoreductase activity, and acyl-CoA oxidase activity (Fig. 4D) while the downregulated genes were enriched in exopeptidase activity, protein metabolic process, positive regulation of lipid transport, organonitrogen compound metabolic process, cytosolic ribosome, and amide metabolic process (supplementary Fig. S2A). The GSEA analysis reveals similar results in GO enrichment. Notable upregulated processes include "U2 type spliceosomal comples" and "Regulation of RNA splicing" while the downregulated processes include “Regulation of cellular response to growth factor stimulus” (Fig. 4E, supplementary Fig. S2E). GSEA plot showing the enrichment of the U2-type spliceosomal complex (GO:0005684 CC) in the ranked gene list (Fig. 4F). The red line represents the enrichment score (ES), which peaks at 0.7178, indicating significant enrichment. The nominal P-value is 0, and the false discovery rate (FDR) is 0.0023, confirming the statistical significance. The normalized ES is 2.3618, which further highlights the significant up-regulation of this term. The KEGG analysis showed a similar result, which the upregulated genes were enriched in spliceosome, replication and repair, protein export, peroxisome, and DNA replication (supplementary Fig. S2B). The downregulated genes were enriched in multiple metabolic processes, glycan biosynthesis, and protein kinases (supplementary Fig. S2C).

In our study, a significant downregulation was observed in members of the intraflagellar transport (*IFT*) and tubulin tyrosine ligase-like (*TTLL*) families among the differentially expressed genes. Specifically, the expression levels of IFT-A complex proteins (*IFT122* and *IFT43*) and IFT-B complex proteins (*IFT25*, *IFT22*, and *IFT20*) were markedly reduced. Similarly, several *TTLL* family members—including *TTLL2*, *TTLL4*, *TTLL10*, and *TTLL11*—also exhibited decreased expression (Fig. 4C), suggesting potential impairment in tubulin modification dynamics (Janke et al., 2005). Conversely, we observed upregulation of vesicular transport machinery, including coatomer and adaptin complexes. Furthermore, both the catalytic α subunit (*PRKAA1*) and regulatory β subunits (*PRKAB1*/*PRKAB2*) of AMPK showed concurrent upregulation.

Weighted gene correlation network analysis (WGCNA) was used to find the co-expression genes, co-expression genes were clustered by expression patterns as represented by the dendrogram and correlation heat map. Clusters of like-regulated genes are referred to as modules by color (supplementary Fig. S3A). The co-upregulated genes in cyst were enriched in ubiquitin protein ligase activity, ubiquitin-protein transferase activity, regulation of cell migration, and endosome (supplementary Fig. S3B). The co-downregulated genes in cyst were enriched in lyase activity, cell cycle G2/M phase transition and calcium−dependent protein kinase (CDPK) activity (supplementary Fig. S3C). The correlation functional networks were used to identify the hub genes in cyst, unfortunately, few hub genes were annotated in our analysis.

### Proteome analysis identifying key proteins and functional pathways during cyst formation

In order to elucidate the regulatory mechanisms governing gene expression during the encystment process, we further performed a comprehensive proteomic analysis. PCA analysis was employed to illustrate the correlations among samples, revealing a high degree of similarity within both the cyst and oxy group (Fig. 5A). Similar to the case in transcriptome (Fig. 4B), cells in cyst stage had dramatically shifted proteome profiles from those in vegetative stage (Fig. 5B). In the proteomics data, we identified 451 upregulated genes and 283 downregulated genes during encystment (Fig. 5C). The KEGG pathway enrichment analysis showed that the upregulated proteins were related to proteasome, spliceosome, and exosome (Fig. 5D). The GO enrichment analysis showed the upregulated proteins were enriched in proteasome and its catabolic process, while the downregulated proteins were enriched in lipid metabolic and cellular modified amino acid metabolic processes (Fig. 5E, supplementary Fig. S4).

**Figure 5.**
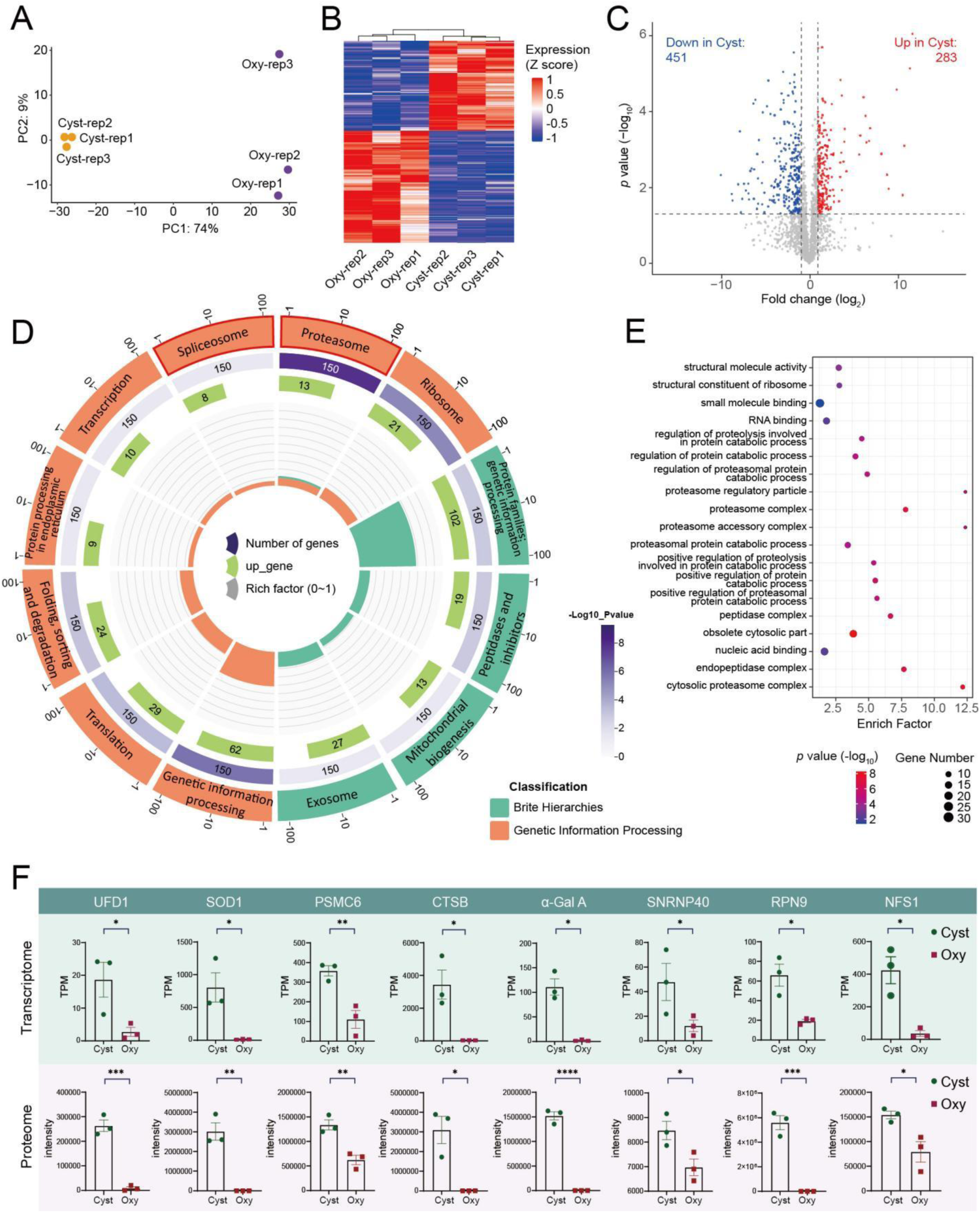
Protein changes between vegetative and cyst stages of *Oxytricha granulifera*. (A) Principal component analysis (PCA) based on proteome profiles. Each point represents a sample. Orange dots represent the vegetative stage, while purple dots represent the cyst stage. (B) Heatmap of proteomic across different stages. (C) Volcano plot of proteomic with fold-change thresholds of 2 and a P-value threshold of 0.05. (D) KEGG circle plot. The inner ring is a bar plot where the height of the bar indicates the enrich factor of the term, the color indicates KEGG terms that the corresponding proteins are associated with. The outer ring displays the upregulated proteins in each term. The third ring shows the total number of proteins associated in this term and color corresponds to the significance (-log10 P-value). The outermost ring displays the names of the KEGG term. The enriched term associated with spliceosome is highlighted in red. (E) GO enrichment. (F) Plots showing RNA and protein (bottom) abundance of key genes. RNA abundance is measured as transcript per million (TPM).

We compared the abundance of seven genes that play important roles in encystment between RNA and protein levels. Transcriptome and proteome analyses confirmed the high expression of these proteins during the encystment period (Fig. 5F). We observed a significant upregulation of UFD1 (Ubiquitin Folding Domain Containing 1), SOD1 (superoxide dismutase 1), PSMC6 (Proteasome 26S Subunit, ATPase, 6), CTSB (Cathepsin B), α-GalA (Alpha-Galactosidase A), SNRNP40 (Small Nuclear Ribonucleoprotein Polypeptide 40), RPN9 (Ribophorin 9) and NFS1 (NAD(P)H:quinone oxidoreductase 1). UFD1, PSMC6, CTSB, RPN9 were involved in the intracellular degradation of proteins, supporting the cell’s transition into the encysted state. SOD1, NFS1 may play a key role in protecting cell from ROS (Alvarez et al., 2017; Fujii et al., 2022). SNRNP40 participates in the biogenesis of small nuclear ribonucleoproteins (snRNPs), potentially influencing the splicing and maturation of pre-mRNA (Will & Lührmann, 2011). α-Gal A, which may be involved in the metabolism of cell surface glycoproteins and glycolipids, affecting cell adhesion and signal transduction (Brockhausen, 2006; Krstić Ristivojević et al., 2020).

### Higher frequency of alternative splicing events and potential roles of 6mA in cyst

In both transcriptome and proteome data, we observed upregulation of several genes or proteins associated with spliceosome, indicating the presence of numerous alternative splicing events during encystment (Fig. 4D, Fig. 5D). Considering the global transcription level difference between encysts and vegetative cells, we analyzed the frequency of alternative splicing events relative to the number of total transcripts and the result indicated that alternative splicing occurred more frequently during encystment (Fig. 6A). Regarding the types of alternative splicing, there were slightly more events during encystment than during the vegetative stage, primarily involving intron compatible, retained introns and other type of alternative splicing (Fig. 6B). We also examined the number of exons per transcript, and the results showed that the majority of transcripts during both stages had one exon (Fig. 6C). However, transcripts with two exons were significantly more common during encystment, possibly reflecting the gene expression characteristics during this stage.

**Figure 6.**
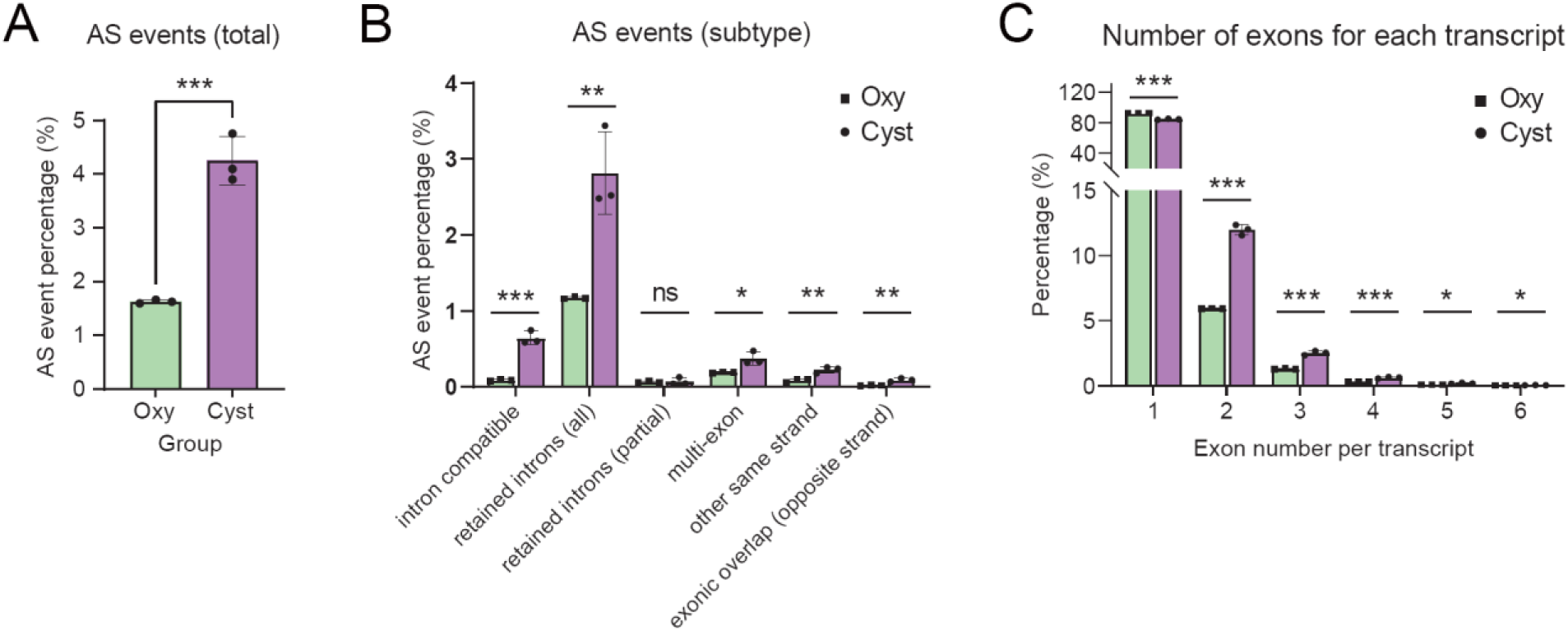
Alternative splicing analysis during the vegetative cell-cyst transition of *Oxytricha granulifera*. (A) Relative frequencies of alternative splicing events compared to the number of total transcripts in cells at vegetative stage (Oxy) and cyst stage (Cyst). (B) Subtype of alternative splicing events between vegetative stage (Oxy) and cyst stage (Cyst). (C) Number of exons for each transcript between two groups. *, p < 0.05; **, p < 0.01; ***, p<0.001.

DNA N6-adenine methylation (6mA) is a transcriptional activator involved in gene regulation and catalyzed by 6mA methyltransferase (6mA MTase). An expanding list of studies shows that 6mA responds to external stress. For example, in the worm *Caenorhabditis elegans*, 6mA is involved in mitochondrial stress adaptation (Ma et al., 2019). In addition, in *Pseudocohnilembus persalinus*, the methyltransferase PpAMT1 regulates cellular growth and encystment by modulating 6mA levels, a conserved regulatory mechanism among ciliates, and reducing the levels of AMT enzymes accelerates encystment, highlighting the critical role of these enzymes in controlling the encystment process (Liu et al., 2024).

In this study, we identified the methyltransferases responsible for 6mA methylation in *O. granulifera*, confirmed the conserved domains of the AMT family using NCBI CD-search (Fig. 7A), and reconstructed the phylogenetic tree of AMT family sequences from representative eukaryotic species (Fig. 7B). Similar to *P. persalinus*, we found that the homologous genes of AMT1 and AMT6/7 in *O. granulifera* are also significantly downregulated during encystment (Fig. 7C). This suggests that 6mA may have a similar negative regulatory role during encystment in *O. granulifera*.

**Figure 7.**
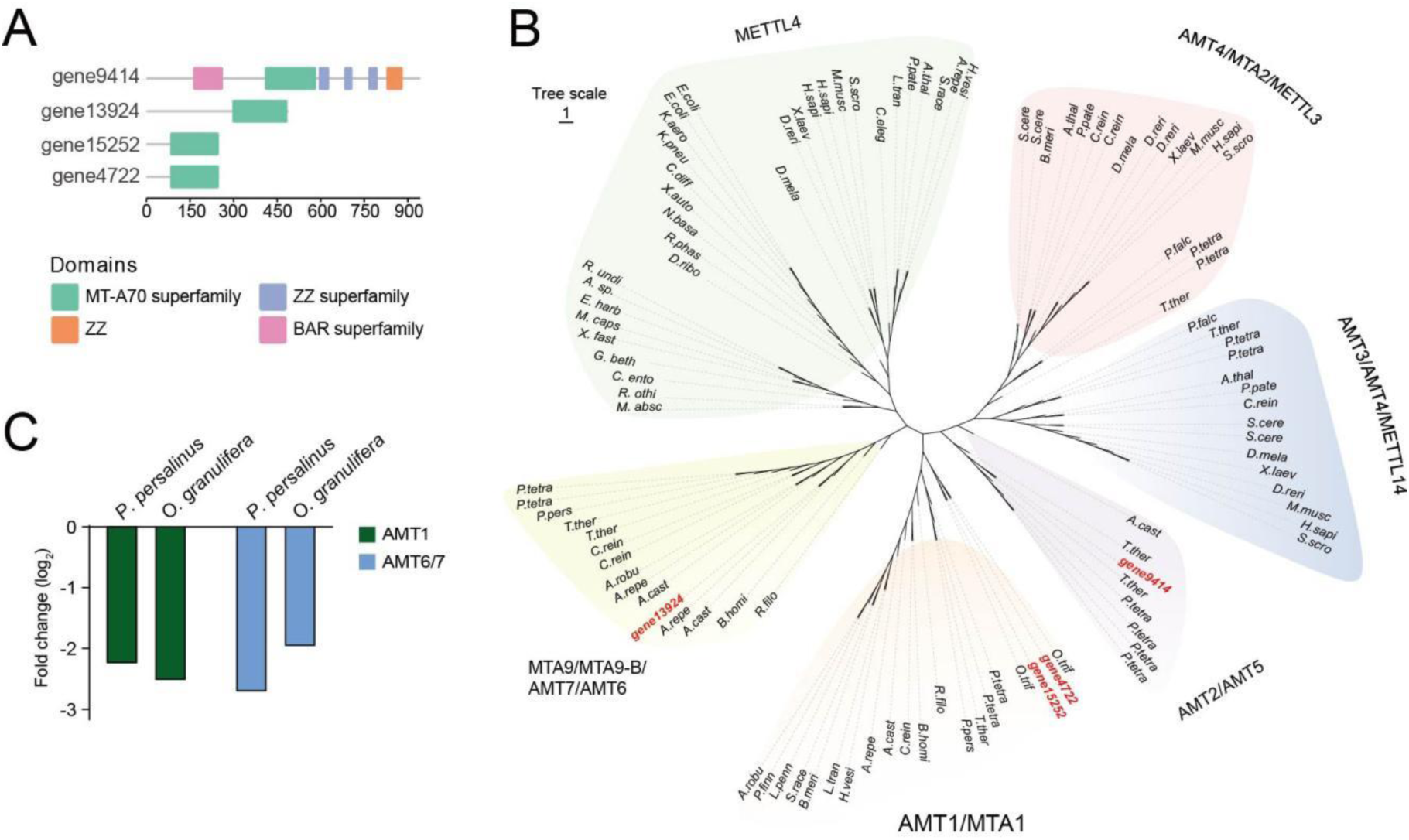
Phylogenetic analysis and domain structure of OgAMT proteins and their differential expression ratios. (A) Domain structures of four potential OgAMT proteins. (B) Phylogenetic analysis of MT-A70 proteins. Full species names are listed in supplementary Table S2. (C) Gene expression levels of potential AMT proteins in *Pseudocohnilembus persalinus* and *Oxytricha granulifera*.

## DISCUSSION

### Evolutionary expansion of gene families provides insights into dormancy regulation

The formation of dormant cysts involving complex and highly regulated physiological and biochemical alterations which might be reflected in the genome evolution (Jin et al., 2024). Through comprehensive analyses of gene family expansion of *Oxytricha granulifera*, we discuss the possible integrated stress response network and organelle-specific adaptations coordinately govern encystation.

The initiation of ciliates dormancy involves environmental signal perception and signal transduction, and the elevated intracellular Ca^2+^ concentrations play the core role by modulating intracellular signaling pathways (Jiang et al., 2018; Pan et al., 2019; Yamaoka et al., 2004). In *O. granulifera*, we identified several expanded gene families that are significantly enriched in functions related to environmental stress response (e.g., “cellular response to environmental stimulus” and “response to water deprivation”), suggesting an enhanced capacity for environmental stress perception in this species. Correspondingly, the enrichment of "response to calcium ion" indicates a possible reinforced regulatory capacity of Ca^2+^ signaling pathways. Additionally, the enrichment of “monoatomic ion transport” indicates the enhancement of the overall capacity of re-establishing cellular ion homeostasis throughout the encystation process, other than the rapid ion-mediated responses in the initial stage.

In ciliates, extrusomes exhibit diverse fates within cysts, distinct from their roles during the trophic phase. For instance, in *Colpoda cucullus* and *Tetrahymena rostrata*, mucocysts expel mucus to form the cyst wall (Funatani et al., 2010; Zebrun et al., 1967; McArdle et al., 1980). In *Pseudourostyla cristata,* protrichocysts not only contribute to the cyst wall but also undergo autophagy (Zhang et al., 2011; Grim, 1985). In *Oxytricha granulifera*, the presence of autophagic vesicles engulfing mucocysts and glycoproteins within cysts suggests that mucocysts are degraded via autophagy rather than secreted. Cyst wall of ciliates is rich of glycoproteins in composition (Izquierdo et al., 2000). We speculate that the significant expansion of gene families related to carbohydrate metabolism (e.g., “hydrolase activity”) may enable *O. granulifera* to utilize the glycoproteins in extrusome components not only as a sustained energy source during early encystment but also as potential materials for cyst wall constituents. This highlights the multifaceted utilization of expanded genes and specialized organelles by the ciliate.

Functioning as a crucial cellular energy sensor, AMPK exerts inhibitory effects on various anabolic pathways, including fatty acid metabolism and glycolysis/gluconeogenesis, thereby facilitating cellular survival during the encystment phase (Jiang et al., 2018; Jung et al., 2010). The coordinated enrichment of “AMP-activated protein kinase activity” and “negative regulation of TORC1 signaling” suggests a potential enhancement of AMPK’s capacity to activate autophagy, possibly through suppressing TORC1-mediated inhibition. Previous studies have shown that cyst cells maintain a low respiration rate, with mitochondria undergoing degradation or aggregation and exhibiting minimal activity during encystment (Funatani et al., 2010). Most notably, the expansion of mitophagy-related genes and macroautophagy aligns with the frequent encapsulation of mitochondria within autophagic vacuoles (Fig. 1J, M). This selective mitochondrial clearance likely serves multiple critical functions: reducing oxidative phosphorylation activity to minimize energy consumption and recycling cellular components to sustain prolonged metabolic quiescence.

In the dormant forms of other groups of single-celled eukaryotes, such as yeast spores (*Saccharomyces cerevisiae*) and resting cells of diatoms (e.g., *Thalassiosira pseudonana*), reduced intracellular pH is a characteristic feature of encystment and this acidification contributes to the preservation of the native structure of proteins and their oligomeric states, as observed in yeast spores (Munder et al., 2016; Wang et al., 2024; Petrovska et al., 2014). The enrichment of “intracellular pH reduction” and “vacuolar acidification” in gene expansion of *O. granulifera* may be associated with enhanced autophagy-mediated waste degradation, as well as reduced metabolism while maintaining the structure and function of important proteins.

### Multilayered regulation of encystment

#### Signal transduction regulation

The AMPK is a highly conserved heterotrimeric kinase complex composed of a catalytic (ɑ) subunit and two regulatory (β and γ) subunits (Ross et al., 2016). The *PRKAA* encodes the α-subunit of AMPK, serving as the direct executor of AMPK kinase activity and a central regulator of cellular energy homeostasis, autophagy, and stress responses. Under energy stress, AMPKα phosphorylates ULK1 at Ser317/Ser777, triggering autophagosome formation, while concurrently alleviating mTORC1-mediated suppression of ULK1 (Ser757 phosphorylation) (Gwinn et al., 2008; Kim et al., 2011). In contrast, the *PRKAB* (encoding the β-subunit) primarily stabilizes the AMPK heterotrimeric complex and modulates its activation through glycogen-binding capacity and subcellular compartmentalization (McBride et al., 2009; Xiao et al., 2013). During *O. granulifera*’s encystation, bioenergetic stress induces transcriptional upregulation of *PRKAB*, leading to AMPK complex activation. The activated AMPK through *PRKAA1*, phosphorylates downstream effectors to suppress mTOR signaling, thereby modulating autophagic flux.

#### Gene expression regulation

In *P. persalinus*, the dynamic 6mA modification exhibits a transition from symmetric to asymmetric sites during the vegetative cell-to-cyst transformation, which is closely associated with gene expression regulation, and the reduction in 6mA levels facilitates cyst formation (Liu et al., 2024). CDPKs are the most abundant class of calcium sensors, being found in protozoa, ciliates, and plants (Harper & Harmon, 2005). Emerging evidence suggests their potential functional importance in *Cryptosporidium parvum* (an apicomplexa) growth (Zhang et al., 2020). Since both CDPK and 6mA-related enzyme genes were downregulated, we propose that this suppression inhibits vegetative growth while promoting cyst formation, with 6mA reduction potentially serving as a key inducer in *O. granulifera*.

#### RNA processing regulation

Alternative splicing (AS) is the process through which primary transcripts can be modified in different arrangements to produce functionally distinct mature mRNAs, and AS events are regulated to ensure production of appropriate protein isoforms in the correct cellular environments (Jacob et al. 2017). Splicing was carried out in a stepwise coordinated fashion by a large ribonucleoprotein complex, named spliceosome—a molecular machine composed of five ribonucleoprotein particles (snRNPs: U1, U2, U4, U5 and U6) (Wahl et al., 2009). Our data show upregulation of the U2 snRNP spliceosome and U5 snRNP in transcriptomic GO enrichment, along with the spliceosome pathway in proteomic KEGG enrichment. Additionally, the upregulation of *U2AF* (U2 snRNP component for splice site recognition) and *SNRNP40* (core protein component of U5 snRNP) among differentially expressed genes (DEGs) suggests heightened spliceosomal activity during encystment formation. This coordinated induction of both early (U2-associated) and late (U5-associated) splicing factors may facilitate proteome remodeling through enhanced RNA processing, potentially regulating stage-specific physiological adaptations required for encystment. Alternative splicing is commonly classified into seven types of simple binary events, among which intron retention refers to an intron remaining in the mature mRNA instead of being spliced out (Jacob & Smith, 2017). Intron retention is the most prevalent mode of alternative splicing in unicellular eukaryotes, and it may lead to alterations in the secondary structure of mRNA, thereby affecting ribosome binding and translation initiation, and ultimately reducing protein synthesis (Grabski et al., 2021; Grau-Bove et al., 2018; McGuire et al., 2008; Monteuuis et al., 2019). Intron retention increases during cyst formation of *Acanthamoeba castellanii* (Amoebozoa), indicating a potential mechanism of gene regulation that could help downregulate metabolism (de Obeso Fernandez Del Valle et al., 2022; Ner-Gaon et al., 2004). In the study on yeast, intron retention has been linked to cellular responses to starvation and stress (Morgan et al., 2019; Parenteau et al., 2019). The enrichment of alternative splicing events among upregulated genes in *O. granulifera* and a high frequency of intron retention and intron compatible alternative splicing suggest that intron retention is likely a common mechanism for single unicellular eukaryotes to under stressful conditions.

### Survival strategy of morphological transformation to reduce energy costs

Morphologically, the encystment is characterized by the formation of cyst-specific structures and the dedifferentiation of vegetative motility and feeding structures. Oxytrichids usually have four layers (ectocyst, mesocyst, endocyst and granular layer) in cyst wall (Gutierréz et al. 2003). In our TEM observation, only two layers cyst wall were found in the encysting cyst, and a large number of cortical electron dense granules which are the same size of the precursors of granular layer in other oxytrichids (Gutiérrez, 1983). Concurrently, expression of coatomer and adaptin related genes was concurrently upregulated. Coatomer constitutes the structural protein core of COPI complex. The COPI complex transports vesicles backward from the Golgi to the ER and within the Golgi, keeping organelles balanced. Adaptin complexes cooperate with clathrin to mediate endocytic and late secretory transport (Field & Dacks, 2009; Robinson, 2004). In *Colpoda steinii*, the cyst-wall precursors—ER-derived small vesicles abundant during encystment but absent from vegetative cells—originate exclusively from the rough endoplasmic reticulum (Ruthmann & Kuck, 1985). It was thus speculated that upregulated vesicular transport-related genes is likely important for cyst wall formation by orchestrating the vesicular transport of the precursors.

Ciliature, as the major component of the tubulin cytoskeleton of ciliate cells, undergoes varying degrees of resorption during encystment in different ciliates groups, resulting in three artificially divided types of cysts: non-kinetosome-resorbing cysts (only the ciliary shafts were partially dedifferentiated); partial-kinetosome-resorbing cysts (most ciliature components dedifferentiated, while some kinetosomes remained intact); kinetosome-resorbing cysts (all the cilia, kinetosomes and microtubules were absorbed) (Martín-González, 1992; Walker & Maugel, 1980; Berger, 1999; Li et al., 2022). Given the absence of ciliary components within the cysts, the cysts are of the kinetosome-resorbing type. The observed expansion of gene families associated with "regulation of microtubule cytoskeleton organization" and "cytoskeletal protein binding" suggests enhanced molecular capacity for both complete ciliary structure disassembly during encystment and efficient cytoskeletal reconstitution during excystation.

The downregulation of *TTLL* and *IFT* might help reprograming ciliature during encystment of *O. granulifera*. Ciliary assembly and maintenance are governed by bidirectional intraflagellar transport (IFT), whose two principal subcomplexes—IFT-A and IFT-B—execute distinct cargo itineraries: IFT-A mediates retrograde trafficking of membrane proteins, whereas IFT-B orchestrates anterograde delivery of soluble precursors, including tubulin (Nakayama & Katoh, 2020). The down-regulation of genes encoding the bidirectional IFT machinery indicates that a precisely balanced IFT flux is indispensable for cilium biogenesis, structural homeostasis, and signal transduction. *TTLL* orchestrate tubulin assembly, maintenance, and motility by encoding enzymes that post-translationally modify tubulin via polyglutamylation, polyglycylation, and tyrosination (Janke & Bulinski, 2011; Mukai et al., 2009). In *P. cristata* cyst, the downregulated *TTLL11* lead to the speculation that the poor microtubule transport functions and facilitate the entry of into dormancy (Pan et al. 2021), whereas more genes of TTLL family were identified in *O. granulifera* further confirmed this idea. Although autophagy can dismantle cilia during encystment of ciliates (Li et al., 2022), our findings indicate that transcriptional fine-tuning of the cilia-dynamic related genes, rather than wholesale degradation, is the primary mechanism preserving ciliary architecture in the cyst.

Downregulation of cytoskeleton related genes and proteins induces structural alterations in cells, reduces cellular volume and water content, which leads to the stagnation of related essential physiological processes such as locomotion and feeding behaviors. At the same time, our results demonstrate that pathways related to Metabolism, Glycan biosynthesis, and Carbohydrate metabolism are significantly downregulated in the KEGG enriched pathways (supplementary Fig S1C). These findings indicate a metabolic shift toward energy conservation during the encystment process in *O. granulifera*.

### Protein degradation, autophagy and homeostasis maintaining

The formation of cysts and the regression of certain cellular structures require the involvement of autophagy and protein degradation pathways. The ubiquitin-proteasome system (UPS) is a crucial pathway for intracellular protein degradation, marking proteins for degradation through ubiquitination and then degrading them by the proteasome (Kwon & Ciechanover, 2017; Schreiber & Peter, 2014). In our study, we observed an enrichment of the ubiquitin-related protein degradation pathway in both the transcriptome and proteome (e.g., UFD1, PSMC6, CTSB and RPN9) (Fig. 5F). The expression of ubiquitin cascade components E2, E3 in ubiquitin degradation signaling pathway was significantly upregulated, suggesting that the UPS is highly activated, which responds to the fast and efficient degradation of damaged and unneeded cellular proteins. PSMC6, also known as proteasome 26S subunit, ATPase 6, is a component of the 19S regulatory particle of the 26S proteasome, which plays a crucial role in the degradation of proteins marked with ubiquitin (Pan et al., 2019; Schreiber & Peter, 2014; Sujashvili, 2008). The unnecessary cellular proteins are degraded which meet the needs of the cell morphological upheaval and rapid cell shrinkage during the cyst formation. This mechanism is indispensable for the encystment process of ciliates, enabling their survival and adaptation to dynamic environmental challenges (Pan et al., 2019).

Autophagy is a process by which cellular proteins and damaged or excessive organelles are degraded through the formation of a double-membrane structure known as the autophagosome (Mizushima, 2007; Yang & Klionsky, 2010). The mTOR complex is an important regulatory molecule that inhibits autophagy, and its negative regulation promotes autophagy (Jung et al., 2010; Yu et al., 2010). Transcriptomic GO enrichment analysis revealed downregulated mTOR signaling and upregulated proteasome-related pathways, consistent with the observation of autophagic vacuoles encapsulating organelles and supported by the expansion of autophagy-related gene families. This multi-layered evidence highlights autophagy as a central mechanism for ciliate encystment, safeguarding cellular survival through efficient resource recycling and organelle quality control in a metabolically quiescent state.

Several oxidoreductase activity proteins, including Hsp70 and GPX8, were identified as regulators during encystment, primarily through their role in scavenging organic peroxides (Pan et al., 2020; Xu et al., 2023). Our findings demonstrate significant upregulation of antioxidant proteins SOD1 and NFS1 in both transcriptomic and proteomic profiles of cysts, along with marked transcriptional elevation of GPX8 (Fig. 5F). This suggests their synergistic role in protecting cells from oxidative stress through enhanced oxidoreductase activity. This enhanced antioxidant defense works synergistically with metabolic reprogramming, structural remodeling, and protein homeostasis to establish a new steady state during the transition from vegetative growth to encystment.

### Conclusions

In our study, we comprehensively investigated the encystment of *O. granulifera*, obtaining detailed morphological changes, assembling a relatively complete macronuclear sequence for the first time, and acquiring both transcriptomic and proteomic data across the vegetative and cyst stages. At the cellular structural level, *O. granulifera* undergoes significant changes characterized by the dedifferentiation of ciliature and mucocysts, as well as the formation of autophagic vacuoles and a cyst wall. At the molecular level, calcium-related genes function as molecular sensors and, in conjunction with 6mA, mediate the initiation of early encystment. The downregulation of genes involved in ciliogenesis and ciliary transport is closely associated with these morphological changes. Meanwhile, the autophagy pathway and the ubiquitin-proteasome system (UPS) are activated, and the expression of stress-responsive proteins, including Hsp70 and GPX8, is upregulated. These changes facilitate the breakdown and reuse of cellular materials, enhancing cellular stress adaptation and homeostasis. Additionally, the upregulation of alternative splicing, particularly in the intron retention pattern, serves as a regulatory mechanism for gene expression associated with low metabolism, thereby increasing the functional diversity of proteins. These findings provide a comprehensive understanding of the encystment process in ciliates, highlighting the coordinated changes at both morphological and molecular levels (Fig. 8).

**Figure 8.**
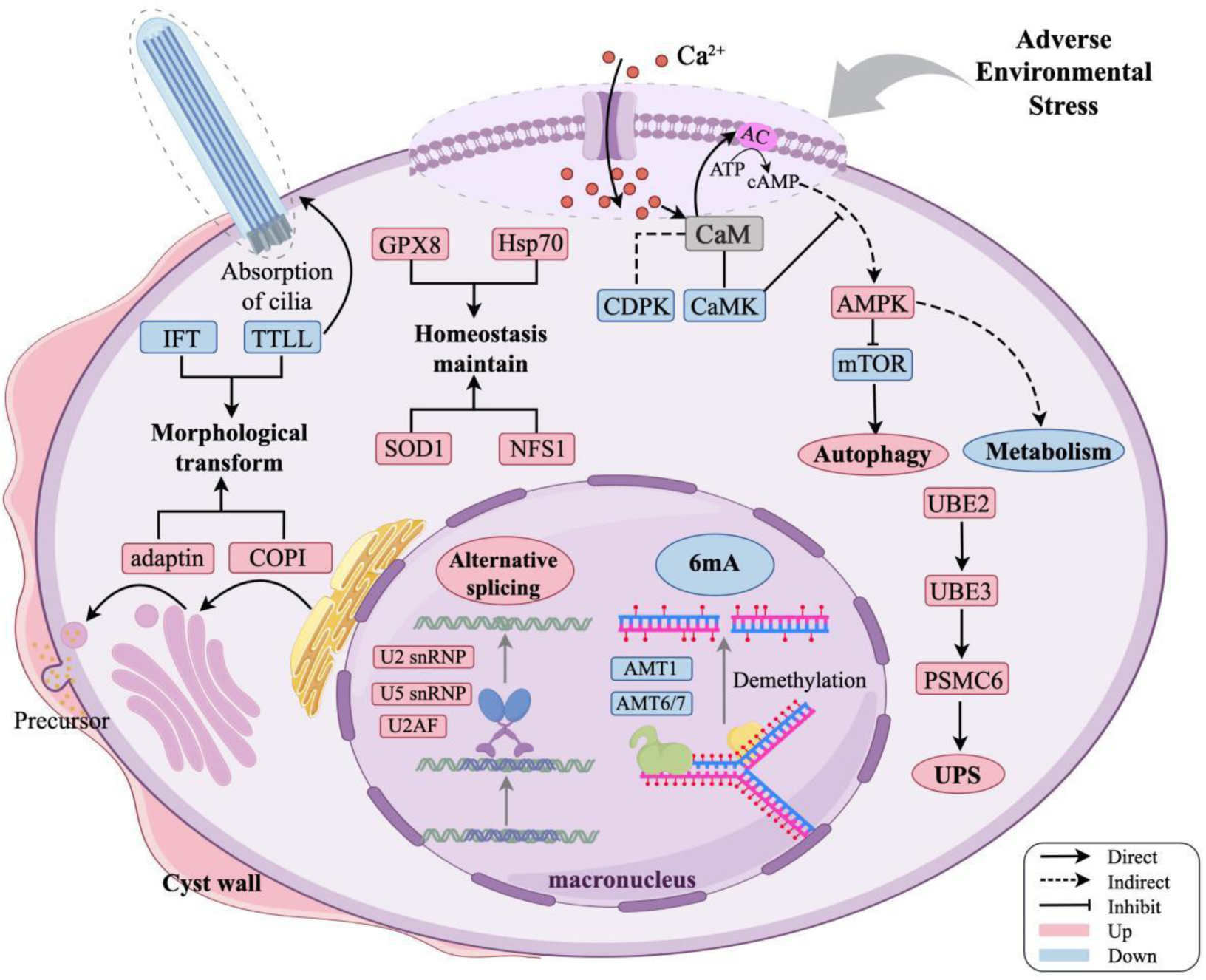
Schematic diagram of major morphological changes and hypothetical regulating signaling network during encystment of *Oxytricha granulifera.* Pink and blue colors indicate upregulation and downregulation of molecular events.

## Author Contributions

**Designed research:** Xinpeng Fan, Xiao Chen, Miaomiao Wang and Juan Yang; **Samples collection:** Tian Wang and Zina Lin; **Molecular experiment:** Tao Hu and Tian Wang; **Morphological observation:** Tao Hu and Zina Lin; **Bioinformatic analyses:** Juan Yang; Visualisation: Juan Yang and Miaomiao Wang; **Statistical analyses:** Zijia Liu, Juan Yang and Miaomiao Wang; **Supervision:** Juan Yang, Miaomiao Wang, Xiao Chen and Xinpeng Fan; **Writing – original draft:** Juan Yang and Miaomiao Wang; **Writing – reviewing and editing:** Juan Yang, Miaomiao Wang, Xiao Chen and Xinpeng Fan; **Funding acquisition:** Xinpeng Fan and Xiao Chen.

## Acknowledgments

This work was supported by the National Natural Science Foundation of China (32170446, 32270512) and Youth Innovation Team of Shandong Provincial Higher Education Institutions.

## Data availability

The genomic reads, and RNA-seq reads have been deposited in the GenBank with accession number PRJNA1302648. The final genome assembly have been deposited into CNSA (https://db.cngb.org/cnsa/) with accession number CNP0007892.

